# Fascin regulates protrusions and delamination to mediate invasive, collective cell migration *in vivo*

**DOI:** 10.1101/734475

**Authors:** Maureen C. Lamb, Kelsey K. Anliker, Tina L. Tootle

## Abstract

Fascin is an actin bundling protein that is essential for developmental cell migrations and promotes cancer metastasis. In addition to bundling actin, Fascin has several actin-independent roles. Border cell migration during *Drosophila* oogenesis provides an excellent model to study Fascin’s various roles during invasive, collective cell migration. Border cell migration requires Fascin. Fascin functions not only within the migrating border cells, but also within the nurse cells, the substrate for this migration. Loss of Fascin results in increased, shorter and mislocalized protrusions during migration. Data supports the model that Fascin promotes the activity of Enabled, an actin elongating factor, to regulate migration. Additionally, loss of Fascin inhibits border cell delamination. These defects are partially due to altered E-cadherin localization in the border cells; this is predicted to be an actin-independent role of Fascin. Overall, Fascin is essential for multiple aspects of this invasive, collective cell migration, and functions in both actin-dependent and -independent manners. These findings have implications beyond *Drosophila*, as border cell migration has emerged as a model to study mechanisms mediating cancer metastasis.

## Introduction

Fascin is an actin-binding protein that bundles or cross-links actin filaments (Hashimoto et al., 2011; Jayo and Parsons, 2010) to promote cell motility and invasion through the formation of filopodia and invadopodia (Adams, 2004; Li et al., 2010; Zanet et al., 2012). While Fascin does promote cell migration in this actin-dependent manner, novel actin-independent roles of Fascin have been discovered (Anilkumar et al., 2003; Jayo et al., 2016; Villari et al., 2015). Fascin directly binds the Linker of the Nucleoskeleton and Cytoskeleton (LINC) complex, which mediates mechanotransduction. Perturbing this interaction impairs nuclear shape deformation essential for single-cell invasive migration (Jayo et al., 2016). Fascin also binds to microtubules and loss of this interaction increases the stability of cellular adhesions causing slower migration (Villari et al., 2015). Additionally, Fascin interacts with Protein Kinase C (PKC), LIM kinases (LIMKs), and, notably, Enabled (Ena; (Anilkumar et al., 2003; Hashimoto et al., 2007; Jayo et al., 2012; Winkelman et al., 2014). Ena is an actin elongation factor, and *in vitro* Ena processivity is increased on Fascin-bundled actin (Harker et al., 2019; Winkelman et al., 2014). These studies illustrate Fascin has multiple functions within the cell that regulate cell migration.

Fascin is important for both developmental cell migrations and cancer metastasis (Cohan et al., 2001; Hashimoto et al., 2011; Hashimoto et al., 2005). Fascin controls cell migration during development such as, growth cone extension, dendrite formation, and in embryonic fibroblasts (De Arcangelis et al., 2004; Ma et al., 2013). Fascin is also highly upregulated in certain types of cancer, and elevated expression is associated with increased invasiveness, aggressiveness and mortality (Arlt et al., 2019; Hashimoto et al., 2011; Hashimoto et al., 2007). While Fascin has been studied in the contexts of 2D migration and single cell 3D migration, the roles of Fascin in invasive, collective cell migration have yet to be investigated (Adams, 2004; Jayo et al., 2016).

*Drosophila* oogenesis – specifically border cell migration – is an ideal model to study invasive, collective cell migration. *Drosophila* oogenesis has 14 developmental stages of the egg chambers or follicles (Spradling, 1993). Each follicle is composed of a single oocyte, 15 germline-derived nurse cells, and a layer of somatic epithelial cells, or follicle cells, surrounding the outside. During Stage 9 (S9) of follicle development, a group of follicle cells at the anterior end are specified to become border cells. This group of 8-10 border cells delaminates from the epithelium and migrate between the nurse cells to reach the nurse cell-oocyte boundary (Montell, 2003; Montell et al., 2012). Delamination is a highly regulated process in which the border cells must maintain cellular adhesions, such as E-cadherin, amongst themselves, but sever adhesions with their neighboring follicle cells and nurse cells (Cai et al., 2014; Montell, 2003). Additionally, border cell migration is very dynamic with protrusions extending and retracting to move the cluster (Fulga and Rorth, 2002; Prasad and Montell, 2007). Upon completing its migration, the border cells produce the micropyle, the structure through which sperm fertilize the egg (Montell et al., 1992; Spradling, 1993). Importantly, the migrating border cells highly express Fascin (Cant et al., 1994). Therefore, we can study the role of Fascin in invasive, collective cell migration *in vivo* using the simple and genetically tractable model of border cell.

Here we find that Fascin plays a critical role in regulating border cell migration. Using a new quantification method that assesses border cell migration during S9, we find loss of Fascin results in significant delays in border cell migration. Surprisingly, Fascin is necessary in both the germline cells and somatic cells but is only sufficient within the somatic cells to promote border cell migration. Live imaging reveals that loss of Fascin results in border cell clusters with more protrusions emerging from all sides that are shorter in length and duration. These alterations culminate in the *fascin*-null clusters migrating slower than controls. These defects are due, in part, to Fascin’s role in regulating Ena. Dominant genetic interactions reveal Fascin and Ena work together to regulate border cell migration, and overexpression of Ena suppresses migration defects in *fascin*-null mutants. Fascin also regulates border cell delamination. In *fascin*-null mutants, the clusters take longer to delaminate, and display altered localization of E-cadherin. Overall, our data reveal that Fascin regulates multiple aspects of border cell migration, including both protrusion dynamics and delamination. These findings suggest that Fascin regulates invasive, collective cell migration through modulating cellular protrusions by bundling actin and regulating cellular adhesions to control initiation of migration.

## Results

### Fascin is required for border cell migration

Previously, it was reported that loss of Fascin does not affect border cell migration (Cant et al., 1994). This analysis showed that border cells of *fascin*-null mutants completed their migration between the nurse cells and reached the nurse cell-oocyte boundary by Stage 10A (S10A) (Cant et al., 1994); we have reproduced these findings (Fig. S1). These findings are surprising as Fascin is highly expressed in the border cells (Cant et al., 1994) and is known to regulate many types of cell migration (Hashimoto et al., 2011; Jayo and Parsons, 2010). Thus, we hypothesized that while border cells reach the nurse cell-oocyte boundary by S10, border cell migration may be delayed during S9 in *fascin* mutants.

Border cell migration is a highly regulated event; at any point during S9, the distance the border cells have migrated is approximately equal to the distance the outer follicle cells are from the anterior (Fig 1A). Thus, delayed or accelerated migration of the border cells can be quantitatively assessed by comparing their location relative to that of the outer follicle cells. Specifically, by subtracting the distance of the outer follicle cells from the anterior end of the follicle from the distance the border cells have migrated we can calculate the migration index (Fig. 1A). A migration index of ∼0 indicates on-time migration, while a negative value indicates delayed migration and a positive value indicates accelerated migration (Fig. 1A).

**Figure 1:**
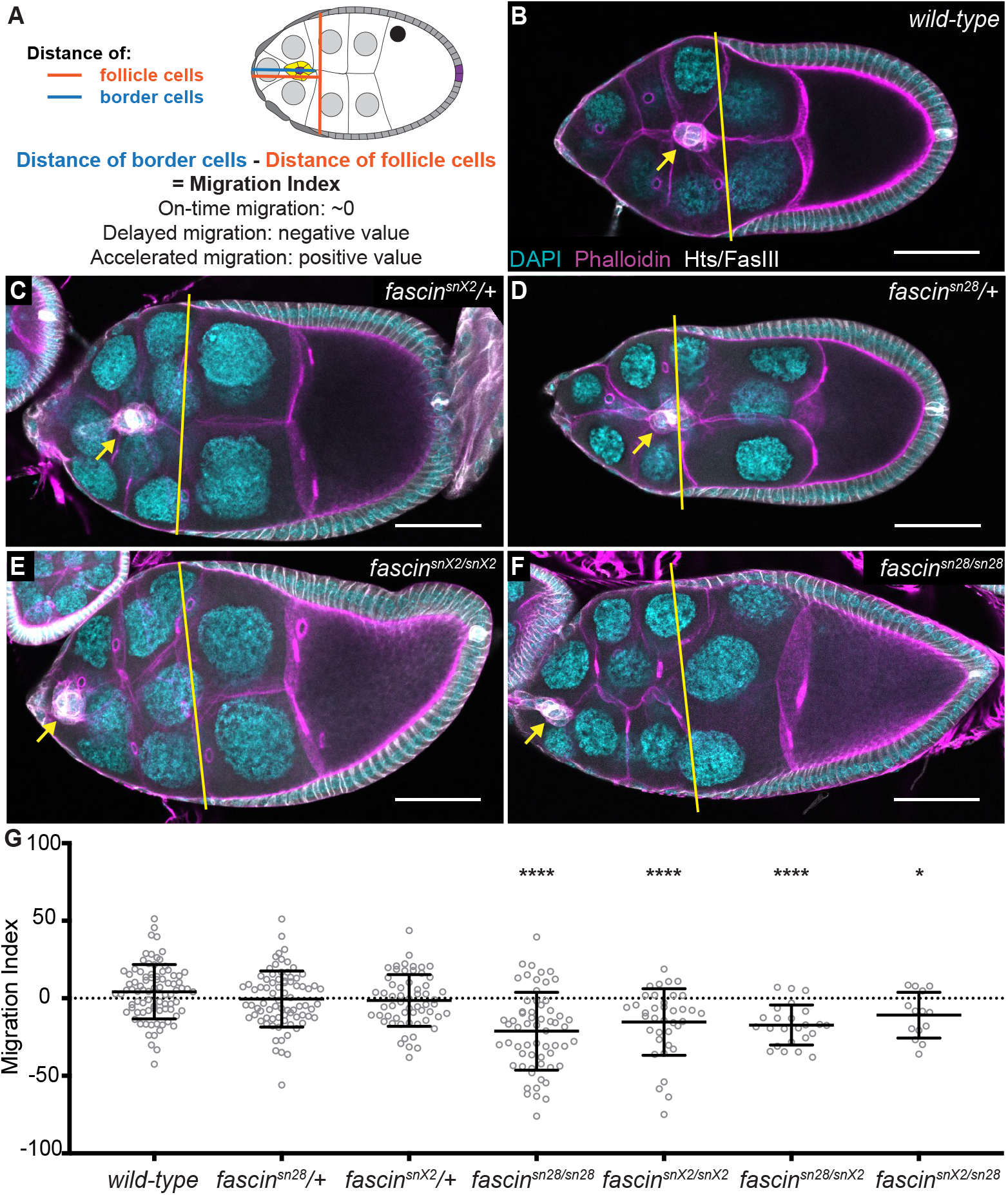
Fascin is required for border cell migration. (**A**) Schematic of migration index quantification for border cell migration during S9. The migration index is the distance of the outer follicle cells from the anterior end of the follicle subtracted from the distance the border cell cluster has migrated. A value of ∼0 indicates on-time migration, negative values indicate delayed migration and positive values indicate accelerated migration. (**B-F**) Maximum projections of 2-4 confocal slices of S9 follicles of the indicated genotypes. Merged images: Hts/FasIII (white, border cell migration stain), phalloidin (magenta), and DAPI (cyan). Yellow lines denote the distance the outer follicle cells have traveled and yellow arrows denote the border cell cluster. Black boxes added behind text to improve text clarity. Scale bars = 50μm. (**B**) wild-type (*yw*). (**C**) *fascin^snX2^*/+. (**D**) *fascin^sn28^*/+. (**E**) *fascin^snX2/X2^*. (**F**) *fascin^sn28/sn28^*. (**G**) Migration index quantification of the indicated genotypes. Dotted line at 0 indicates on-time migration. Each circle represents a single S9 follicle. *p<0.01, ****p<0.0001. Loss of Fascin results in significant border cell migration delays during S9 (E-F) compared to the heterozygous *fascin* mutants (C, D) and wild-type (B) controls using the migration index quantification of border cell migration (G).

To assess border cell migration during S9, we performed immunofluorescent staining for Hts and FasIII, this stain labels both border cells (yellow arrow) and outer follicle cells (yellow line) and enables us to assess border cell migration and quantify migration index (Fig. 1B-G). This stain will be referred to throughout the paper as the border cell migration stain. Using this, we quantified migration index in wild-type and *fascin* mutant follicles (Fig. 1B-G). Two different null alleles of *fascin* were used, *fascin^sn28^* and *fascin^snX2^* (Cant and Cooley, 1996; Cant et al., 1994). S9 follicles that were wild-type or heterozygous for mutations in *fascin* display on-time border cell migration with the border cell cluster being in line with the outer follicle cells (Fig. 1B-D and G; migration indices of 4.23, −0.48 and −1.35, respectively). Loss of Fascin by both homozygous (*fascin^sn28/sn28^* and *fascin^snX2/snX2^*) and transheterozygous *fascin* mutations (*fascin^sn28/snX2^* and *fascin^snX2/sn28^*; maternal allele is listed first) results in border cell clusters that are significantly delayed (Fig. 1E-G; migration indices of −21.20 (p<0.0001), −15.27 (p<0.0001), −17.25 (p<0.0001) and −10.85 (p=0.0363), respectively). These data reveal Fascin is required for proper on-time border cell migration during S9 of *Drosophila* oogenesis.

### Fascin is necessary in both the germline and somatic cells for border cell migration

We next sought to identify where Fascin is needed for border cell migration. During S9, while Fascin is most highly expressed in the border cell cluster, the nurse cells and outer follicle cells also express Fascin (see Fig. S2A; (Cant et al., 1994)). Moreover, the nurse cells are the substrate upon which the border cells migrate and changes in nurse cell structure or stiffness perturb border cell migration (Aranjuez et al., 2016; Cai et al., 2016).

We used the UAS/GAL4 system (Rorth, 1998) to express Fascin RNAi constructs to knockdown Fascin in specific cell types and determine the effects on border cell migration. Two different Fascin RNAi lines were used (second chromosome: TRiP.HMJ21813 and third chromosome: TRiP.HMS02450) and yielded similar results; data presented uses the third chromosome line. We knocked down Fascin in all somatic cells (*c355* GAL4), in the border cells (*c306* GAL4), or in the germline cells (*matα* GAL4). Knockdown of Fascin was confirmed by immunostaining for Fascin (Fig. S2A-E’). Knockdown of Fascin in all somatic cells (c355 GAL4) causes signification border cell migration delays compared to the GAL4 driver only and RNAi only controls (Fig. 2A, B, E; migration indices of −22.97 compared to 0.76 and −7.14; p<0.0001). Similarly, knockdown of Fascin in only the border cells (*c306* GAL4) causes delayed migration, however, this is not significant compared to controls (Fig. 2A, C, E; migration indices of −13.94 compared to −4.92 and −7.14; p=0.051). This mild phenotype is likely due to insufficient knockdown of Fascin during migration, as high levels of Fascin in observed in the border cells at early stages of migration (Fig. S2D-D’) and diminishing levels at the later stages (Fig. S2E-E’). Notably, knockdown of Fascin in the germline cells (*matα* GAL4) also significantly delays border cell migration compared to controls (Fig. 2A, D, E; migration indices of −27.80 compared to −7.25 and −7.14; p<0.0001). These findings indicate Fascin is necessary in both the somatic and germline for border cell migration.

**Figure 2:**
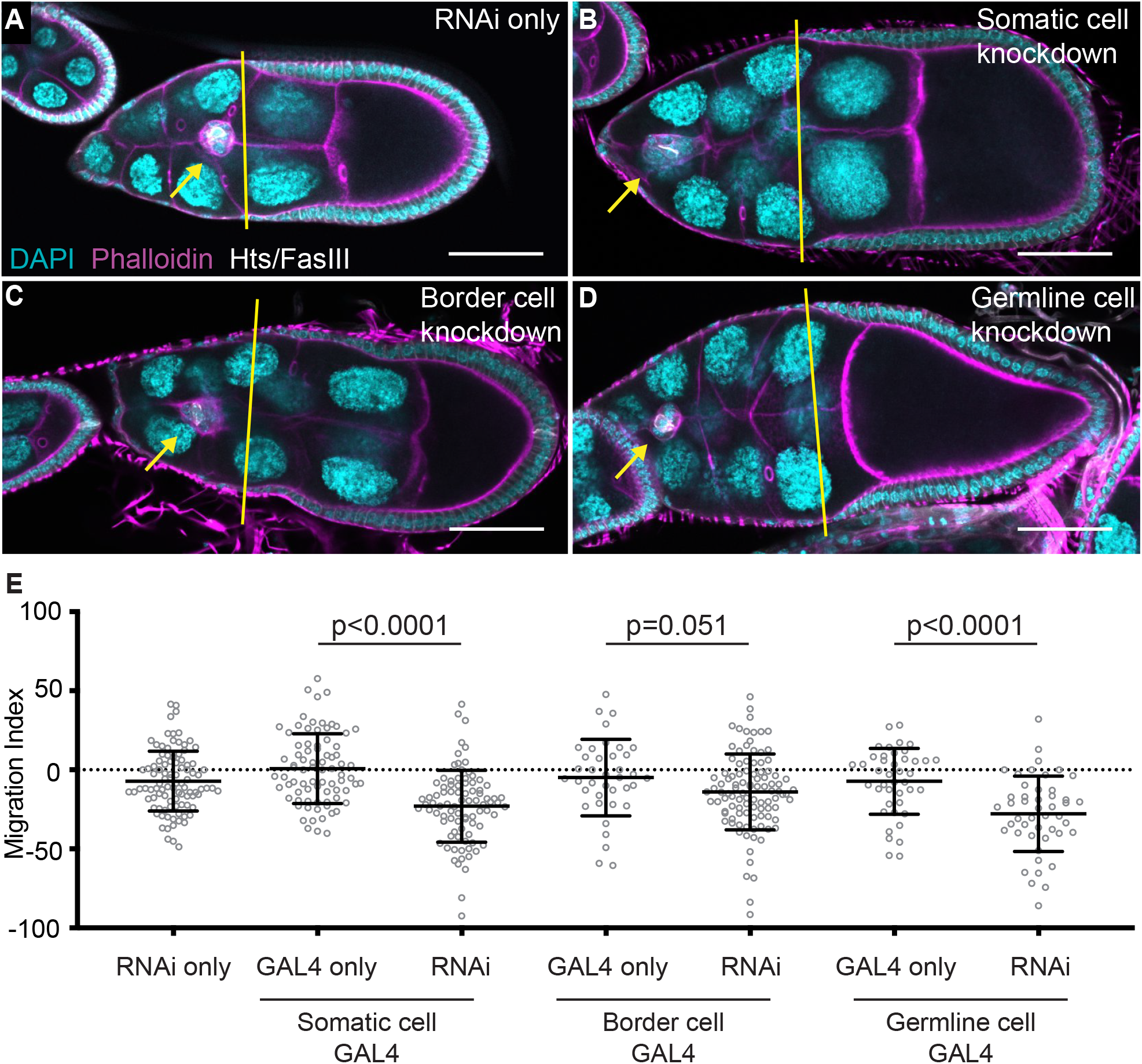
Fascin is necessary in both the germline and somatic cells for border cell migration. (**A-D**) Maximum projections of 2-4 confocal slices of S9 follicles of the indicated genotypes. Merged images: Hts/FasIII (white, border cell migration stain), phalloidin (magenta), and DAPI (cyan). Yellow lines denote the distance the outer follicle cells have traveled and yellow arrows denote the border cell cluster. Black boxes added behind text to improve text clarity. Scale bars = 50μm. (**A**) RNAi only (*fascin RNAi*/+). (**B**) Somatic cell knockdown of Fascin (*c355 GAL4/+; +/fascin RNAi)*. (**C**) Border cell knockdown of Fascin (*c306 GAL4*/+; +/*fascin RNAi*). (**D**) Germline cell knockdown of Fascin (*matα GAL4(3)/fascin RNAi*). (**E**) Migration index quantification of the indicated genotypes. Dotted line at 0 indicates on-time migration. Each circle represents a single S9 follicle. Fascin is necessary for border cell migration in both the somatic and germline cells of the follicle. Border cell migration is significantly delayed in follicles with somatic (B, E) and germline knockdown of Fascin (D, E). Knockdown of Fascin in only the border cells does not cause significant border cell migration delays (C, E); this is likely due to poor knockdown efficiency of the GAL4 driver (see Fig. S2).

### Somatic expression of Fascin rescues border cell migration

We next asked in what cell types is Fascin sufficient for normal border cell migration. The UAS/GAL4 system was used to express GFP-Fascin in specific cell types of *fascin* mutant follicles to determine where restoring expression rescues border cell migration. We expressed GFP-Fascin in the somatic cells (c355 GAL4), the germline cells (*oskar* GAL4), or in both the germline and somatic cells (*actin5C* GAL4) (see Table S1 for all statistical comparisons). Expression of GFP-Fascin in the somatic cells of *fascin* mutant follicles restores border cell migration (Fig. 3A, B, G; migration indices −0.35 compared to −20.19; p=0.0004). Conversely, expression of GFP-Fascin in the germline cells of *fascin* mutant follicles fails to rescue border cell migration (Fig. 3C, D, G; migration indices of −20.97 compared to −26.49; p=0.49). Finally, expression of Fascin in both the somatic and germline cells of *fascin* mutant follicles restores border cell migration (Fig. 3E-G; migration indices of 10.99 compared to −26.43; p<0.0001). Thus, Fascin is necessary but not sufficient in the germline cells, whereas Fascin is both necessary and sufficient within the somatic cells to promote proper border cell migration.

**Figure 3:**
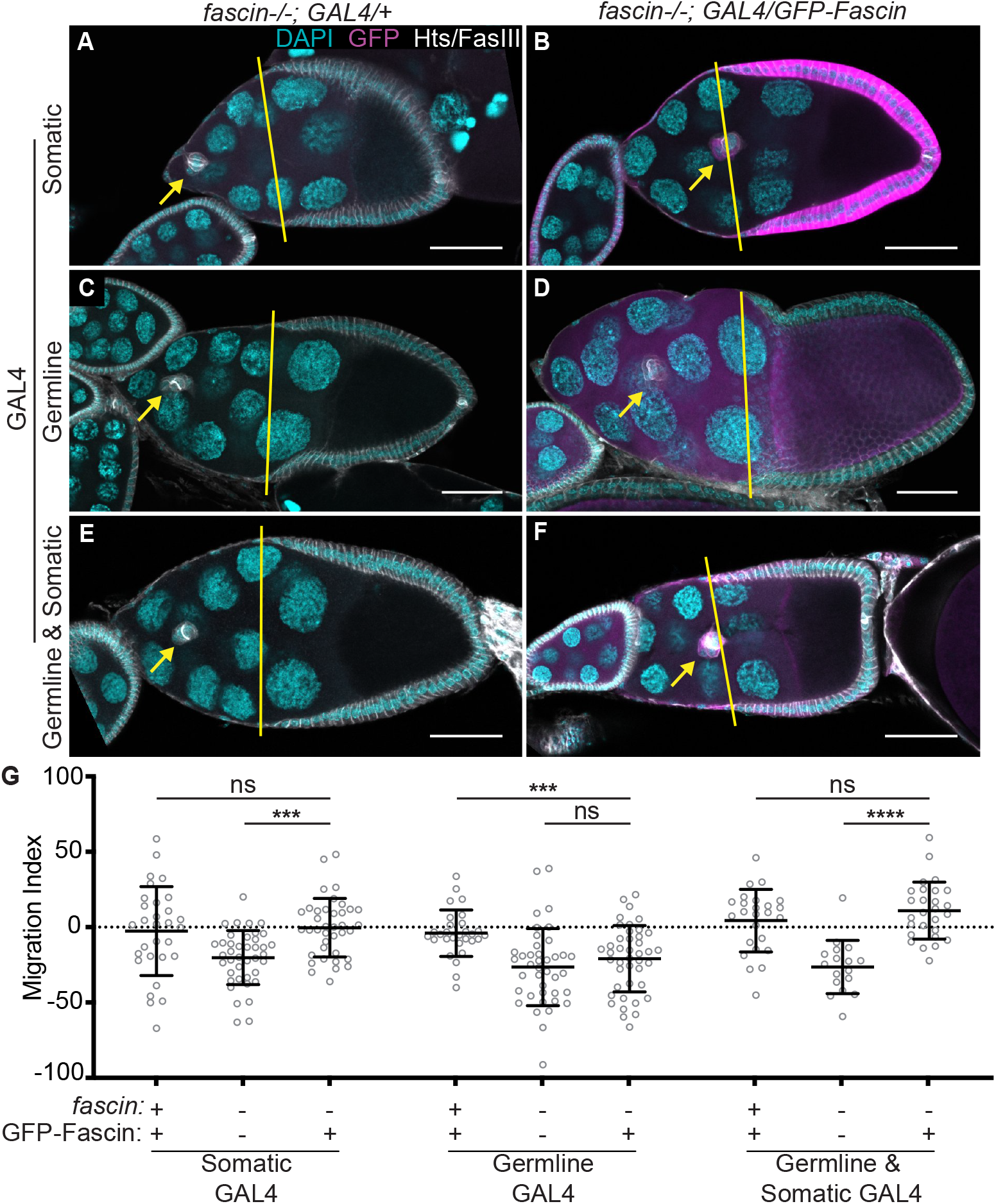
Somatic expression of Fascin rescues border cell migration. (**A-F**) Maximum projections of 2-4 confocal slices of S9 follicles of the indicated genotypes. Merged images: Hts/FasIII (white, border cell migration stain), GFP (magenta), and DAPI (cyan). Yellow lines denote the distance the outer follicle cells have traveled and yellow arrows denote the border cell cluster. Black boxes added behind text to improve text clarity. Scale bars = 50μm. (**A**) *fascin* mutant control with somatic GAL4 (*c355 GAL4, fascin^sn28^/fascin^sn28^*). (**B**) Somatic expression of GFP-Fascin in *fascin* mutant (*c355 GAL4, fascin^sn28/sn28^;* +/*UAS-GFP-Fascin*). (**C**) *fascin* mutant control with germline GAL4 (*fascin^sn28/sn28^; oskar GAL4(2)*/+). (**D**) Germline expression of GFP-Fascin in *fascin* mutant (*fascin^sn28/sn28^; oskar GAL4(2)/UAS-GFP-Fascin*). (**E**) *fascin* mutant control with germline and somatic GAL4 (*fascin^sn28/sn28^; actin5C GAL4/*+). (**F**) Germline and somatic cell expression of GFP-Fascin in *fascin* mutant (*fascin^sn28/sn28^; actin5C GAL4/UAS-GFP-Fascin*). (G) Migration index quantification of the indicated genotypes. GFP-tagged Fascin expression in the *wild-type* background was also included for each GAL4. The ‘+’ and ‘−’ marks for *fascin* underneath the graph indicate whether the follicle was wild-type or *fascin* mutant, respectively. The ‘+’ and ‘−’ marks for GFP-Fascin indicate whether or not the follicle expressed GFP-tagged Fascin under the control of the denoted GAL4 drivers. Dotted line at 0 indicates on-time migration. Each circle represents a single S9 follicle. ns indicates p>0.05, ***p<0.001, and ****p<0.0001. Restoring expression of Fascin in the somatic cells (A, B, G) or in both the somatic and germline cells (E-G) of *fascin*-null follicles rescues border cell migration. Conversely, germline expression of Fascin in *fascin*-null follicles fails to rescue border cell migration (C, D, G).

### Fascin regulates protrusion dynamics in the migrating border cell cluster

To determine how loss of Fascin causes delayed border cell migration we utilized live imaging. We visualized border cell migration with membrane localized GFP expressed under the control of the *slbo* promoter (*slbo>mCD8-GFP*), which specifically labels the border cells and allows us to analyze cluster protrusions.

During migration, the border cell cluster typically forms one or two large protrusions that extend and retract from the leading end of the cluster as it migrates (Bianco et al., 2007; Prasad and Montell, 2007). In agreement with this, control clusters (*fascin^sn28^*/+) typically have one or two main protrusions extending and retracting from the front of the cluster (Fig. 4A-A”, Movie 1). Conversely, in *fascin*-null mutants (*fascin^sn28/sn28^*) the clusters extend many protrusions from their front, sides, and back (Fig. 4B-B”, Movie 2). Clusters in control follicles have just one protrusion in 64% of the frames analyzed versus 34% of the frames in *fascin*-null follicles (Fig. 4C). Furthermore, the clusters in *fascin*-null follicles have a higher percentage of frames with 3-4 protrusions (19%) compared to those of controls (1%) (Fig. 4C; p<0.0001, Pearson’s chi-squared test). Moreover, we assessed the localization of the protrusions on the cluster: front (0° to 45° and 0° to 315°), sides (45° to 135° and 225° to 315°), or back (135° to 225°) of the cluster (Sawant et al., 2018). The *fascin*-null clusters have a significantly altered protrusion localization with 43% of the protrusions emerging from either the side or back of the cluster compared to 17% for the control clusters (Fig. 4D; p<0.0001, Pearson’s chi-squared test).

**Figure 4:**
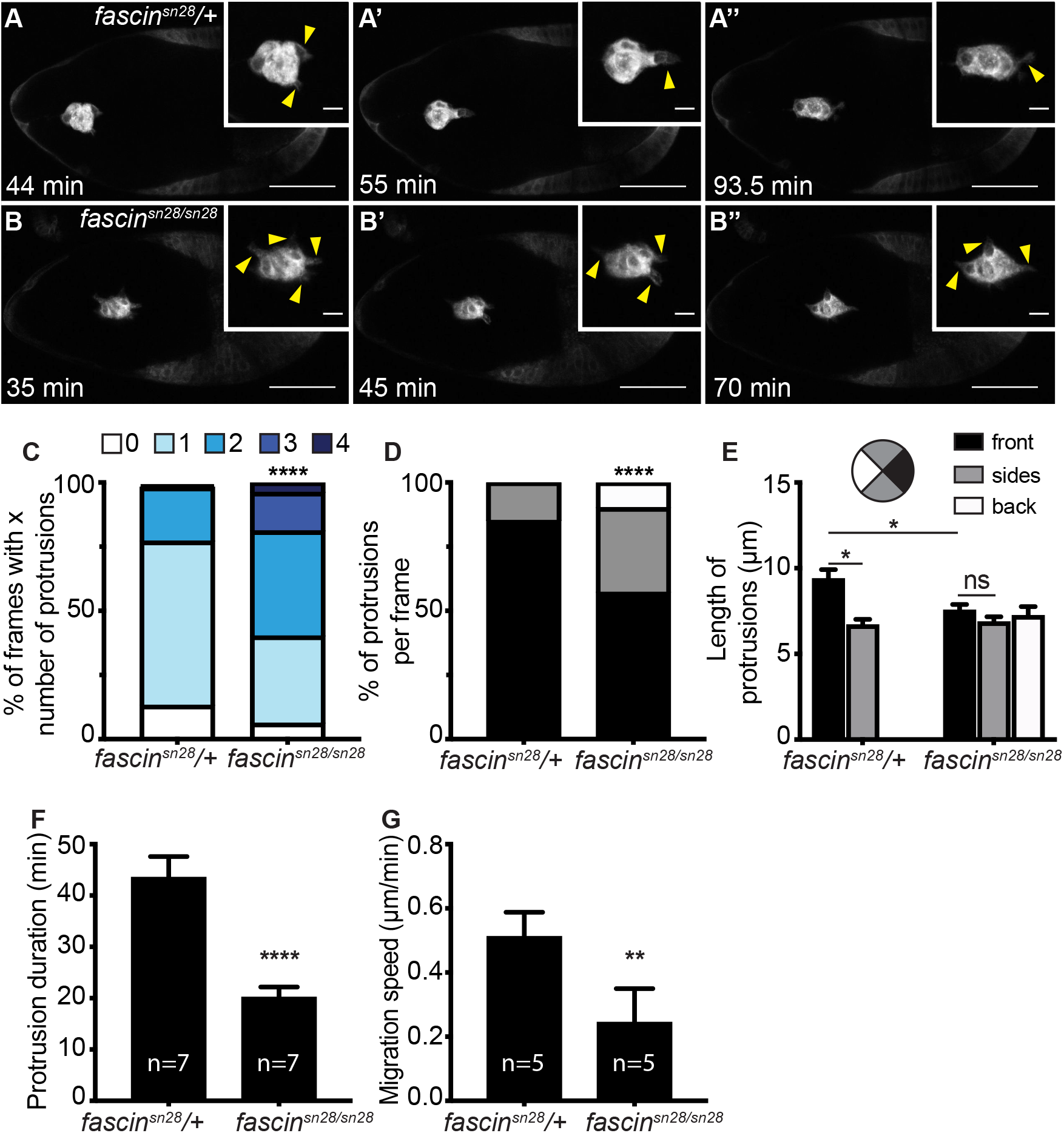
Fascin regulates protrusion dynamics during border cell migration. (**A-B”**) Maximum projection of 2-4 confocal slices from time-lapse live imaging. The border cell cluster was visualized using *slbo>mCD8-GFP* expression and direction of migration is to the right in each image. Insets are zoom-ins of the same border cell cluster and yellow arrowheads indicate protrusions. Time is denoted in minutes (min). Scale bars =50μm for primary images and 10μm for insets. (**A-A”**) control follicle (*fascin^sn28^*/+; Movie 1). (**B-B”**) *fascin*-null follicle (*fascin^sn28/sn28^*; Movie 2). (C-F) Graphs of protrusion dynamics are from control (n=7) and *fascin*-null follicles (n=7). (C) Quantification of the number of protrusions emerging from the cluster per frame. The total number of protrusions per frame was counted for the same number of frames in each video and binned into groups based on total number of protrusions: 0, 1, 2, 3, or 4. ****p<0.0001, Pearson’s chi-squared test. (**D-E**) Quantification of the percent of protrusions per frame (D) and protrusion length (E) based on location of the protrusion. Briefly, protrusions were binned into groups based on if they emerged from the front (0° to 45° and 0° to 315°, black), sides (45° to 135° and 225° to 315°, grey), or back (135° to 225°, white) of the cluster. In D, the same number of frames was analyzed in each video and only frames with at least 1 protrusion were counted; ****p<0.0001, Pearson’s chi-squared test. In E, a protrusion was defined as an extension greater than 4μm long; ns indicates p>0.05, *p<0.05, One-way ANOVA. (**F**) Quantification of protrusion duration. Protrusion duration was defined as the total time elapsed between the protrusion beginning to extend and fully retracting. ****p<0.0001, Student’s t-test. (**G**) Quantification of migration speed. Migration speed was quantified by measuring cluster displacement over time during mid-migration. n=5 for control follicles and n=5 for *fascin*-null follicles. **p<0.01, Student’s t-test. Loss of Fascin results in border cells clusters that have more protrusions (C) that are mislocalized on the cluster (D) and significantly shorter in length (E) and duration (F) compared to control clusters. These changes in protrusion dynamics cause slower migration speeds (G).

In addition to quantifying protrusions per frame, we measured the protrusion length and binned them based on their directionality in the same manner as described above. The protrusions that emerge from the front of the cluster are typically longest in length (Bianco et al., 2007; Prasad and Montell, 2007). Protrusions extending from the front of the cluster were significantly longer in control clusters compared to *fascin*-null clusters (Fig. 4E; 9.3μm compared to 7.5μm, respectively; p=0.045). Additionally, in control clusters, the protrusions extending from the front are significantly longer than the protrusions extending from the sides (Fig. 4E; front=9.3μm, sides=6.6μm; p=0.047). Conversely, *fascin*-null clusters extend protrusions of similar lengths from all sides of the cluster (Fig. 4E; front=7.5μm, sides=6.8μm, and back=7.2μm). Additionally, protrusion duration is significantly shorter in the *fascin*-null clusters, with the average duration being 20min compared to 43.4min for controls (Fig. 4F, p<0.0001).

Lastly, we quantified the migration speed of clusters during mid-migration. Loss of Fascin results in significantly slower migration (0.24μm/min) compared to controls (0.51μm/min; Fig. 4G; p=0.0019). Overall, these data indicate the loss of Fascin impairs protrusion formation and regulation within the cluster, and these impairments cause slower migration speeds.

### Fascin regulates Ena to promote border cell migration

We next wanted to determine how Fascin regulates protrusions during border cell migration. Recent findings demonstrate that Fascin cooperates with the actin elongation factor Ena to promote actin polymerization and filament formation *in vitro* by enhancing Ena processivity (Harker et al., 2019; Winkelman et al., 2014). Additionally, loss of Ena causes border cell migration defects (Gates et al., 2009). Based on these data, we hypothesize Fascin regulates border cell cluster protrusions by promoting Ena activity.

To assess if Fascin regulates Ena during border cell migration we used dominant genetic interactions studies. Reduced levels of Fascin (*fascin-*/+) or Ena (*ena-*/+) alone should be sufficient to maintain normal border cell migration. If Fascin and Ena function together to mediate border cell migration, then reduced levels of both (*fascin-*/+; *ena-*/+) will exhibit delayed border cell migration. Partial loss of Fascin (data not shown) or two different *ena* alleles *ena^210^*/+ (Fig. 5A) and *ena^23^*/+ (data not shown) exhibit on-time border cell migration (Fig. 5C; migration indices of 1.29, −0.85, and −0.86, respectively). However, double heterozygotes of *fascin* with either *ena* allele (*fascin-*/+; *ena-*/+) causes significant border cell migration delays (Fig. 5B-C; migration indices of −15.58 (p=0.0015) and −20.25 (p=0.0002)). While the dominant genetic interaction results support our hypothesis that Fascin regulates Ena to control border cell migration, if our hypothesis is correct then overexpression of Ena is predicted to suppress the border cell migration delay observed in *fascin*-null follicles. Indeed, expression of RFP-tagged Ena in *fascin* mutant follicles restores on-time migration (Fig. 5D-F; migration indices of −5.26 compared to −31.63; p=0.0002). These findings support the model that the defects observed in the protrusion dynamics of *fascin* mutant clusters are due, at least in part, to Fascin’s role in regulating Ena to promote actin elongation.

**Figure 5:**
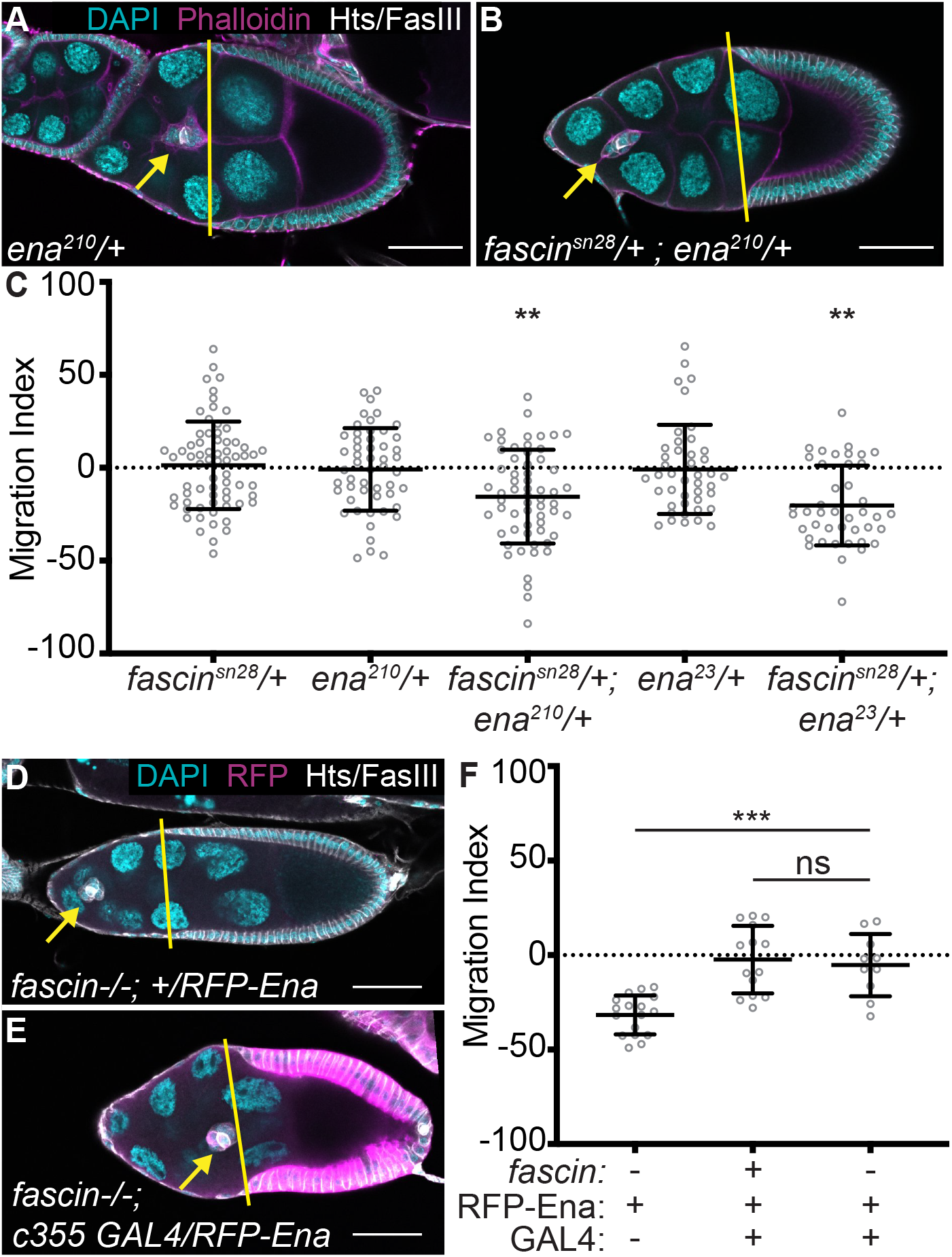
Fascin genetically interacts with Ena to regulate border cell migration. (**A-B, D-E**) Maximum projections of 2-4 confocal slices of S9 follicles of the indicated genotypes. Merged images: (**A-B**) Hts/FasIII (white, border cell stain), phalloidin (magenta), and DAPI (cyan) or (**D-E**) Hts/FasIII (white, border cell stain), RFP (magenta), and DAPI (cyan). Yellow lines denote the distance the outer follicle cells have traveled and yellow arrows denote the border cell cluster. Black boxes added behind text to improve text clarity. Scale bars = 50μm. (**A**) *ena^210^*/+. (**B**) *fascin^sn28^*/+*; ena^210^*/+. (**D**) *fascin* mutant control (*fascin^sn28/sn28^;* +/*UAS-RFP-Ena*). (**E**) Somatic expression of Ena in *fascin* mutant (*c355 GAL4, fascin^sn28/sn28^*; +/*UAS-RFP-Ena*). (**C, F**) Migration index quantification of the indicated genotypes. Dotted line at 0 indicates on-time migration. Each circle represents a single S9 follicle. In F, RFP-tagged Ena expression in the GAL4 background was also included on the graph. The ‘+’ and ‘-’ marks for *fascin* underneath the graph indicate whether the follicle was wild-type or *fascin* mutant, respectively. The ‘ +’ and ‘-’ marks for RFP-Ena indicate expression or not of RFP-tagged Ena by the denoted GAL4 drivers. The ‘ +’ and ‘-’ marks for GAL4 indicate whether the follicle had the somatic GAL4 or not. ns indicates p>0.05, **p<0.01, ***p<0.001 (Student’s t-test). Double heterozygotes for mutations in *fascin* and *ena* exhibit significant delays in border cell migration (A-C). Overexpression of Ena in the somatic cells rescues border cell migration in *fascin*-null follicles (D-F).

### Fascin regulates the delamination of the border cells

In addition to regulating protrusions during migration, Fascin also contributes to border cell delamination. Delamination is the process by which the border cell cluster detaches from the surrounding follicle cells to begin its migration. Live-imaging of follicles during delamination revealed *fascin*-null follicles spend more time detaching from the follicular epithelium (Fig. 6B-B” and Movie 4 compared to 6A-A” and Movie 3). We quantified this change in delamination time by measuring the amount of time elapsed from cluster formation to when the cluster is fully delaminated during early S9. The *fascin*-null clusters take significantly longer to delaminate (320min) compared to control clusters (147min, Fig. 6C; p<0.0001). Additionally, 3 *fascin*-null clusters failed to delaminate during the course of imaging (Fig 6C, indicated by x’s). These results suggest that Fascin promotes border cell delamination.

**Figure 6:**
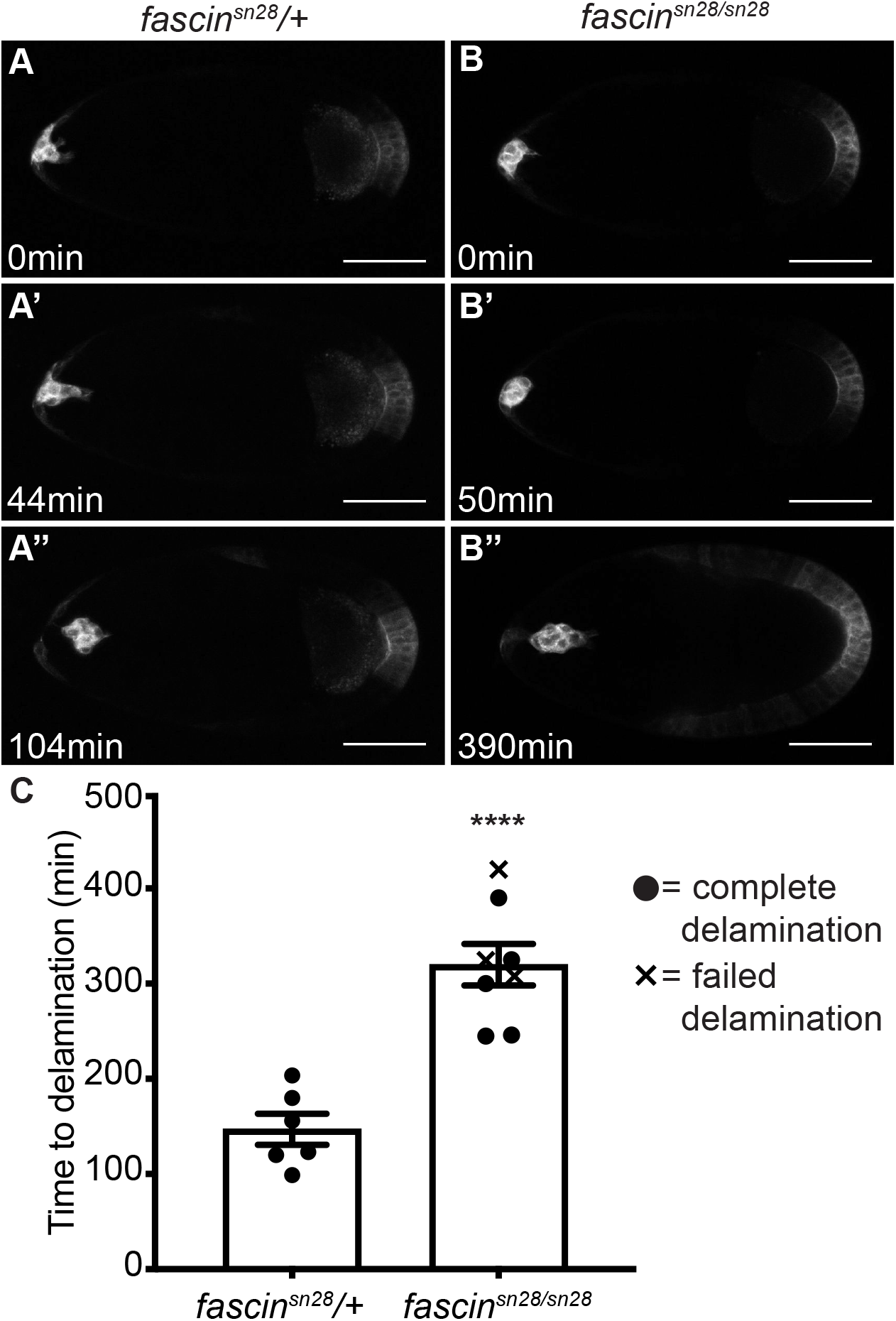
Fascin regulates border cell delamination. (**A-B”**) Maximum projection of 3 confocal slices from time-lapse live imaging. The border cell cluster was visualized using *slbo>mCD8-GFP* expression and direction of migration is to the right in each image. Time is denoted as minutes (min). Scale bars =50μm. (**A-A”**) Control follicle (*fascin^sn28^*/+; Movie 3). (**B-B”**) *fascin*-null follicle (*fascin^sn28/sn28^;* Movie 4). (C) Quantification of time to delamination for control (*fascin^sn28^/+*) and *fascin*-null (*fascin^sn28/sn28^*) follicles. Time to delamination was defined as the amount of time elapsed from early S9 to when the border cell cluster completely detached from the epithelium. Closed circles indicate completed delamination, x’s indicate the cluster did not fully delaminated by the time the imaging ended. n=6 for control follicles and n=8 for *fascin* mutant follicles. ****p<0.0001 (Student’s t-test). *fascin* mutant border cells clusters take significantly longer to delaminate (B-C) compared to the control clusters (A, C).

One process that is essential for delamination is disassembly of cell-cell adhesions between border cells and neighboring follicle and nurse cells (Cai et al., 2014; De Graeve et al., 2012; Niewiadomska et al., 1999). One adhesion molecule that must be regulated is E-cadherin (Cai et al., 2014; De Graeve et al., 2012). Both increasing or decreasing E-cadherin levels in the nurse cells impairs border cell migration (Cai et al., 2014). We were unable to assess dominant genetic interactions between *e-cadherin* and *fascin* mutants because heterozygosity for mutations in *e-cadherin* resulted in border cell migration delays (data not shown). Therefore, we assessed E-cadherin by immunofluorescence. As initial differences in E-cadherin between wild-type and *fascin*-null delaminating clusters were subtle, samples were stained in the same tube for further analyses. Delaminating border cell clusters in *fascin*-null follicles retain intense E-cadherin localization at all cell-cell boundaries (Fig. 7C-D compared to A-B), with stronger E-cadherin intensity at the periphery of the cluster (border cell-nurse cell boundary) compared to control clusters as observed by both intensity labeling (Fig. 7D compared to B, yellow arrowheads) and line-scan analysis (Fig. 7F compared to E). Additionally, unlike *fascin*-null clusters (Fig. 7C-D, F), E-cadherin intensity in control clusters is much lower at the cluster periphery (border cell-nurse cell boundary) than the border cell-polar cell boundaries (Fig. 7A-B, E). These results suggest that Fascin is required for reducing E-cadherin at the border cell cluster boundary, which is necessary for delamination.

**Figure 7:**
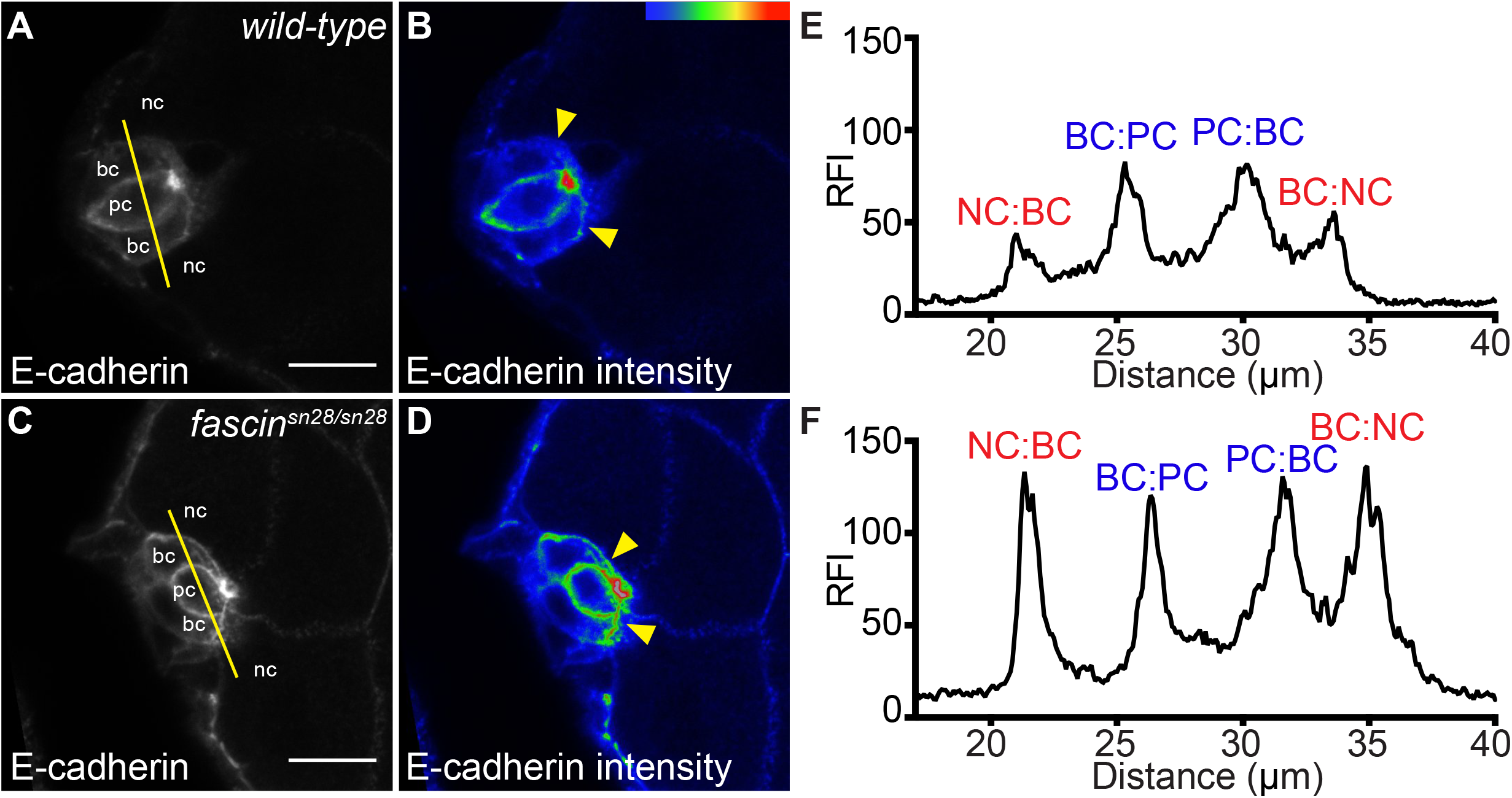
Fascin regulates E-cadherin localization in the delaminating cluster. (**A-D**) Maximum projections of 2 confocal slices of S9 follicles of the indicated genotypes. (**A, C**) E-cadherin (white). (**B, D**) E-cadherin staining pseudocolored with Rainbow RGB, red indicating highest intensity pixels. (**A-B**) wild-type (*yw*) (C-D) *fascin*-null (*fascin^sn28/sn28^*). The cell-types are indicated as nc = nurse cell, bc = border cell, and pc = polar cell. Yellow lines in A, C denote the location where line was drawn for line scan analysis. The yellow arrowheads in B, D denote E-cadherin intensity differences at the border cell-nurse cell boundary. Scale bars = 10μm. (**E-F**) Fluorescence intensity plot of E-cadherin along the yellow lines across border cell cluster (A, C). X-axis: distance; Y-axis: Relative Fluorescence Intensity (RFI). (**E**) Wild-type cluster (*yw*) (**F**) *fascin*-null cluster (*fascin^sn28/sn28^*). NC:BC and BC:NC indicate a nurse cell-border cell boundary (red). PC:BC and BC:PC indicate a polar cell-border cell boundary (blue). The *fascin-* null clusters display overall higher intensity E-cadherin staining compared to wild-type clusters (A-D). The increased E-cadherin localization in the *fascin*-null clusters is most notable at the border cell-nurse cell boundaries (red, F compared to E) compared to the border cell-polar cell boundaries (blue).

## Discussion

Here we provide evidence that Fascin regulates invasive, collective cell migration through multiple functions. Specifically, loss of Fascin results in delays in border cell migration during S9 of *Drosophila* oogenesis (Fig. 1). Fascin functions not only within the border cells, but also in the nurse cells, the substrate on which the border cells migrate, to mediate migration (Figs. 2-3). While Fascin’s role within the nurse cells remains unknown, it may involve both actin-dependent and -independent functions (see further discussion below). Within the border cells, Fascin regulates cluster protrusions (Fig. 4). This regulation is likely achieved through the actin bundling function of Fascin and regulation of the actin elongation activity of Ena (Fig. 5). Additionally, loss of Fascin impairs border cell delamination (Fig. 6), which may be the result of altered E-cadherin localization in the delaminating cluster (Fig. 7). This defect, as discussed below, is likely due to an actin-independent function of Fascin. Ultimately, our findings suggest that Fascin’s roles in invasive, collective cell migration can be attributed to both its actin-dependent and actin-independent functions.

Multiple cell types within the follicle require Fascin for border cell migration. We discovered that Fascin is necessary in both the somatic and germline cells, and sufficient within the somatic cells for border cell migration (Figs. 2-3). The roles of Fascin within the border cells are discussed below, here we speculate on the roles of Fascin within the nurse cells, the substrate on which the border cells migrate. We hypothesize that loss of Fascin alters the stiffness of the nurse cells which impairs border cell migration. Increasing nurse cell stiffness by enhancing non-muscle myosin II contractility impairs border cell migration (Aranjuez et al., 2016; Cai et al., 2016). Interestingly, *in vitro* Fascin inhibits non-muscle myosin II (Elkhatib et al., 2014). Therefore, loss of Fascin in the nurse cells may increase non-muscle myosin II contractility resulting in stiffer nurse cells and delayed border cell migration. Another means by which Fascin may alter the nurse cell stiffness is by controlling nurse cell-nurse cell adhesion. Indeed, we find that E-cadherin levels are higher on all cell membranes, including the nurse cells during S9 (Fig. 7). Such increased adhesion may impede border cell migration. Finally, Fascin may regulate the structure of the cortical actin in the nurse cells to control stiffness, as loss of Fascin results in cortical actin breakdown during mid-oogenesis (Groen et al., 2012). Further studies are needed to understand how Fascin functions within the germline to modulate border cell migration.

One way by which Fascin functions within the border cells to regulate migration is through controlling protrusion formation and dynamics. Loss of Fascin results in shorter and more protrusions (Fig. 4B-E). Consistent with this finding, in both *Drosophila* and cancer cells loss of Fascin results in shorter protrusions during single cell migration (Alam et al., 2012; Zanet et al., 2012). Additionally, Fascin is regulated by and interacts with PKC (Adams et al., 1999) and disruption of this interaction increases cellular protrusions (Anilkumar et al., 2003). Moreover, atypical PKC zeta regulates border cell migration (Wang et al., 2018), but it is unclear if other forms of PKC also do this. Further exploration of PKC regulation of Fascin in border cell protrusion formation and migration is warranted. Protrusion duration is also shorter in *fascin*-null clusters (Fig. 4F). This observation is consistent with the finding that Fascin contributes to protrusion persistence by stabilizing actin bundles (Bear et al., 2000). Fascin may stabilize protrusions by regulating the actin elongation factor Ena. Previous *in vitro* studies uncovered that Ena has increased processivity on actin bundled specifically by Fascin (Harker et al., 2019; Winkelman et al., 2014). These findings suggest that Fascin may regulate the activity of Ena to promote proper protrusion formation and dynamics required for border cell migration. Supporting this idea, dominant genetic interaction studies indicate that Fascin and Ena work within the same pathway to regulate border cell migration (Fig. 5A-C). Additionally, overexpression of Ena restores border cell migration in *fascin* mutants (Fig. 5D-F). These findings indicate that loss of Fascin results in decreased Ena activity and supports the model that Fascin acts to increase Ena processivity to promote stable protrusions necessary for mediating border cell migration. Further studies are needed uncover how this interaction influences protrusions during migration *in vivo*.

Another role of Fascin during border cell migration is regulating the delamination of the cluster. Loss of Fascin results in significantly longer delamination times (Fig. 6). Contributing to this delamination defect is the retention of high levels of E-cadherin on the membranes of the cluster, particularly at the border cell-nurse cell boundaries (Fig. 7). Proper levels of E-cadherin between the nurse cells and border cells are necessary for migration, as knockdown or overexpression of E-cadherin in the border cells or nurse cells results in impaired border cell migration (Cai et al., 2014; Niewiadomska et al., 1999). Therefore, persistence of E-cadherin along this boundary may impair border cell delamination. Other cell adhesion proteins are also required for border cell migration and delamination, including integrins (Dinkins et al., 2008; Villari et al., 2015). Integrins are highly dynamic during cell migration, and either increased or decreased stability of these adhesions impedes migration (Delon and Brown, 2007). Interestingly, Fascin has been proposed to promote integrin-based adhesion dynamics through its interaction with microtubules (Villari et al., 2015). Disruption of Fascin binding to microtubules leads to increased integrin adhesion stability resulting in decreased cell migration (Anilkumar et al., 2003; Villari et al., 2015). This data leads us to speculate that Fascin may control integrin dynamics during border cell delamination and migration through interaction with microtubules. Future studies are needed to test the role of Fascin in regulating cell adhesion dynamics required for border cell delamination and migration.

Fascin may also regulate border cell migration by mediating mechanotransduction. A key mediator of mechanotransduction is the LINC complex. The LINC Complex interacts with the cytoskeletal filaments within the cytoplasm and extends into the nucleus where it interacts with the nuclear lamina (Lombardi et al., 2011; Lombardi and Lammerding, 2011). This structure allows transmission force from the outside of the cell to the nucleus and plays a critical function in regulating nuclear shape and position during invasive cell migration (Alam et al., 2015; Harada et al., 2014; Lombardi et al., 2011; Lombardi and Lammerding, 2011). Fascin binds directly to the cytoplasmic part of the LINC complex in both the *Drosophila* ovary and mammalian cultured cells, and disruption of this interaction impairs nuclear deformation required for mammalian single cell invasive migrations (Jayo et al., 2016). As border cell migration is highly invasive, we hypothesize that disrupting the Fascin-LINC complex interaction will lead to defects in border cell nuclear deformation and impair migration.

The different functions of Fascin must by tightly regulated to ensure they are employed properly to mediate migration. One of the ways that Fascin is regulated is through phosphorylation (Adams et al., 1999; Anilkumar et al., 2003; Zanet et al., 2012). PKC phosphorylates Fascin in response to integrin activation (Anilkumar et al., 2003). Following this phosphorylation Fascin cannot bundle actin and binds to PKC to control integrin dynamics (Anilkumar et al., 2003). Future studies are needed to determine if this interaction controls the balance between Fascin bundling actin and promoting integrin dynamics during border cell migration. Additionally, previous work in our lab demonstrated that Fascin is regulated by prostaglandins (PGs) (Groen et al., 2012). PGs are lipid signaling molecules that regulate a wide variety of biological processes, including cytoskeletal dynamics (Bulin et al., 2005; Peppelenbosch et al., 1993; Tamma et al., 2003; Tootle, 2013). We previously showed that PGs regulate actin bundling during *Drosophila* oogenesis through Fascin (Groen et al., 2012). Exactly how PGs regulate Fascin has yet to be determined, however we believe this may occur through regulating Fascin phosphorylation (Groen and Tootle, unpublished data) and localization (Groen, 2015; Jayo et al., 2016). Indeed, loss of PGs prevents Fascin’s localization to the nuclear periphery where it interacts with the LINC Complex (Jayo et al., 2016). Other work in our lab has found that PGs also regulate border cell migration (Fox, Mellentine, and Tootle, manuscript in preparation). As described above, Fascin regulates Ena during border cell migration, and interestingly, PGs regulate Ena localization and activity in the nurse cells (Spracklen et al., 2014); these findings suggest that PGs may regulate the interaction between Fascin and Ena. Thus, PGs may regulate multiple functions of Fascin to control border cell migration. Altogether, many regulatory mechanisms control how Fascin functions during cell migration, and future studies are needed to define the means of regulating Fascin during border cell migration.

Border cell migration has emerged as an excellent model to study cancer metastasis *in vivo*. Border cell migration recapitulates the collective cell migration often seen in cancer metastasis and enables us to study essential aspects of this migration, such as cluster adhesion or polarization (Friedl and Gilmour, 2009; Montell et al., 2012). Fascin’s role in promoting cancer metastasis is well documented in several types of carcinomas (Gross, 2013; Hashimoto et al., 2011). Fascin is not typically expressed in adult epithelial tissue, however elevated expression of Fascin in epithelial cancers has been correlated with increased aggressiveness, mortality, and notably, metastasis (Arlt et al., 2019; Hashimoto et al., 2011; Yoder et al., 2005). In fact, knockdown of Fascin decreases metastasis in a xenograft tumor model of colon cancer (Hashimoto et al., 2007). Here we identified Fascin as a new regulator of border cell migration and find that Fascin influences both protrusion and adhesion dynamics to control this collective invasive migration. Thus, border cell migration is a simplified, *in vivo*, and genetic tractable system to define the actin bundling-dependent and -independent roles of Fascin in regulating invasive, collective cell migration.

## Materials and methods

### Fly stocks

Fly stocks were maintained on cornmeal/agar/yeast food at 21°C, except where noted. Before immunofluorescence and live imaging, flies were fed wet yeast paste daily for 2-4 days. Unless otherwise noted, *yw* was used as the wild-type control. The following stocks were obtained from the Bloomington Stock Center (Bloomington, IN): *sn^X2^, ena^210^, ena^23^, matα* GAL4 (third chromosome), *c355* GAL4, *c306* GAL4, *actin5C* GAL4, and *UASp-RNAi-Fascin* (TRiP.HMS02450 and TRiP.HMJ21813). The *sn28* line was a generous gift form Jennifer Zanet (Université de Toulouse, Toulouse, France; (Zanet et al., 2012), the *oskar* GAL4 line (second chromosome) was a generous gift from Anne Ephrussi (European Molecular Biology Laboratory, Heidelber, Germany; (Telley et al., 2012), the *UASp-GFP-Fascin* wild-type transgenic fly line was a generous gift from Francois Payre (Université de Toulouse, Toulouse, France; (Zanet et al., 2009), the *UASp-RFP-Ena* wild-type transgenic fly line was a generous gift from Mark Peifer (University of North Carolina, Chapel Hill, NC, unpublished) and the slbo>mCD8-GFP transgenic fly line was a generous gift from Xiaobo Wang (French National Centre for Scientific Research, Toulouse, France). Expression of *UASp-RNAi-Fascin* was achieved by crossing to *mata* GAL4, *c355* GAL4, and *c306* GAL4, maintaining crosses at 25°C and progeny at 29°C. The *sn28, c355* GAL4 flies were generated by recombining *sn28* and *c355* GAL4 onto the same chromosome. Briefly, *sn28, c355* GAL4 males were identified by selecting for the *singed* phenotype (marker for *sn28*) and w+ eyes (marker for *c355* GAL4). Recombination was verified by crossing *sn28, c355 GAL4/FM7* flies to *sn28; UASp-GFP-Fascin* and assessing both GFP expression and *singed* phenotype. A similar recombination scheme was performed to generate *sn28, c306* GAL4/FM7 flies. Expression of *UASp-GFP-Fascin* was achieved by crossing to *oskar* GAL4, *c355* GAL4, and *actin5C* GAL4, crosses were maintained at 25°C and progeny at 29°C. Expression of *UASp-RFP-Ena* was achieved by crossing to *c355* GAL4, crosses were maintained at 25°C and progeny at 29°C.

### Immunofluorescence

Whole-mount *Drosophila* ovary samples were dissected into Grace’s insect media and fixed for 10 minutes at room temperature in 4% paraformaldehyde in Grace’s insect media (Lonza, Walkersville, MD or Thermo Fischer Scientific, Waltham, MA). Briefly, samples were blocked using Triton antibody wash (1X phosphate-buffered saline, 0.1% Triton X-100, and 0.1% bovine serum albumin) six times for 10 minutes each. Primary antibodies were diluted with Triton antibody wash and incubated overnight at 4°C. The following primary antibodies were obtained from the Developmental Studies Hybridoma Bank (DSHB) developed under the auspices of the National Institute of Child Health and Human Development and maintained by the Department of Biology, University of Iowa (Iowa City, IA): mouse anti-Hts 1:50 (1B1, Lipshitz, HD; (Zaccai and Lipshitz, 1996), mouse anti-FasIII 1:50 (7G10, Goodman, C; (Patel et al., 1987); mouse anti-Fascin 1:20 (sn7c, Cooley, L; (Cant et al., 1994), rat anti-DCAD2 1:20 (Umemura, T; (Oda et al., 1994). Additionally, the following primary antibody was used: rabbit anti-GFP 1:2000 (pre-absorbed on *yw* ovaries at 1:20 and used at 1:100; Torrey Pines Biolabs, Inc., Secaucus, NJ) and rabbit anti-dsRed 1:300 (Clontech, Mountain View, CA). After 6 washes in Triton antibody wash (10 minutes each), secondary antibodies were incubated overnight at 4°C or for ∼4 hours at room temperature. The following secondary antibodies were used at 1:500: AlexaFluor (AF)488::goat anti-mouse, AF568::goat anti-mouse, AF488::goat anti-rabbit, AF568::goat anti-rabbit (Thermo Fischer Scientific) and AF647::goat anti-mouse and AF488::goat anti-rat (Jackson ImmunoResearch Laboratories, Inc., West Grove, PA). AF647-, rhodamine, or AF568-conjugated phalloidin (Thermo Fischer Scientific) was included with primary and secondary antibodies at a concentration of 1:250. After 6 washes in Triton antibody wash (10 minutes each), 4’,6-diamidino-2-phenylindole (5 mg/ml) staining was performed at a concentration of 1:5000 in 1X PBS for 10 minutes at room temperature. Ovaries were mounted in 1 mg/ml phenylenediamine in 50% glycerol, pH 9 (Platt and Michael, 1983). All experiments were performed a minimum of three independent times.

### Image acquisition and processing

Microscope images of fixed *Drosophila* follicles were obtained using LAS AS SPE Core software on a Leica TCS SPE mounted on a Leica DM2500 using an ACS APO 20x/0.60 IMM CORR -/D objective (Leica Microsystems, Buffalo Grove, IL) or using Zen software on a Zeiss 700 LSM mounted on an Axio Observer.Z1 using a Plan-Apochromat 20x/0.8 working distance (WD) = 0.55 M27 or a EC-Plan-Neo-Fluar 40x/1.3 oil objective (Carl Zeiss Microscopy, Thornwood, NY). Maximum projections (two to four confocal slices), merged images, rotations, and cropping were performed using ImageJ software (Abramoff et al., 2004).

### Quantification of migration index

Quantification of the migration index of border cell migration during S9 was performed on confocal image stacks of follicles stained with anti-Hts and anti-FasIII. Measurements of migration distances were obtained from maximum projections of 2-4 confocal slices of deidentified 20x confocal images using ImageJ software (Abramoff et al., 2004). Briefly, a line segment was drawn from the anterior end of the follicle to the front or posterior of the border cell cluster and the distance measured, this was defined as the distance of border cell migration. Additionally, a line segment was drawn from the anterior end of the follicle to the anterior end of the main-body follicle cells and the distance measured, this was defined as the distance of the follicle cells. The migration index was calculated in Excel (Microsoft, Redmond, WA) by subtracting the follicle cell distance from the border cell distance. Data was compiled, graphs generated, and statistical analysis performed using Prism (GraphPad Software, La Jolla, CA).

### Line scan analysis of E-cadherin

Line scan analysis was performed on maximum projections of 2 confocal slices of a 40x confocal image using ImageJ software (Abramoff et al., 2004). Briefly, a line segment was drawn across a delaminating border cell cluster and the plot profile function was used to generate a fluorescent intensity plot for E-cadherin. Raw data was graphed in Prism (GraphPad Software). The cell boundaries were defined as the peaks in fluorescent intensity.

### Live imaging

Whole ovaries were dissected from flies fed wet yeast past for 2-3 days and maintained at 25°C until the last 16-24 hours when they were moved to 29°C. Genotypes used for live imaging were *sn28/FM7; slbo>mCD8-GFP* and *sn28/sn28; slbo>mCD8-GFP*. Ovaries were dissected in Stage 9 (S9) medium (Prasad *et al*. 2007): Schneider’s medium (Life Technologies), 0.6x penicillin/streptomycin (Life Technologies). 0.2 mg/ml insulin (Sigma-Aldrich, St. Louis, MO), and 15% fetal bovine serum (Atlanta Biologicals, Flowery Branch, GA). S9 follicles were hand dissected and embedded in 1.25% low-melt agarose (IBI Scientific, Peosta, IA) made with S9 media on a coverslip-bottom dish (MatTek, Ashland, MA). Just prior to live imaging, fresh S9 media was added to coverslip-bottom dish. Live imaging was performed with Zen software on a Zeiss 700 LSM mounted on an Axio Observer.Z1 using a Plan-Apochromat 20x/0.8 working distance (WD) = 0.55 M27 (Carl Zeiss Microscopy, Thornwood, NY). Images were acquired every 5-5.5 mins for at least 3 hours. Maximum projections (two to five confocal slices), merge images, rotations, and cropping were performed using ImageJ software (Abramoff et al., 2004). To aid in visualization live imaging videos and stills were brightened by 50% in Photoshop (Adobe, San Jose, CA).

### Quantification of live imaging

Quantification of live imaging videos was based on analyses done in Sawant et al. (Sawant et al., 2018). Analyses were performed in ImageJ (Abramoff et al., 2004) using maximum projection of 2-5 confocal slices time-lapse videos of border cell migration. Parameters quantified include number of protrusions per frame, protrusion length, protrusion duration, and migration speed. For number of protrusions per frame, the number of protrusions emerging from the front (0° to 45° and 0° to 315°), sides (45° to 135° and 225° to 315°), and back (135° to 225°) of the cluster was counted per frame for an hour of migration. For protrusion length, a protrusion was defined as an extension longer than 4 μm from the cluster body. The length of the protrusions was measured and binned into groups based on the direction emerging from cluster: front (0° to 45° and 0° to 315°), sides (45° to 135° and 225° to 315°), and back (135° to 225°). Protrusion duration was measured by quantifying the amount of time elapsed between the protrusion beginning to extend and the protrusion fully retracting. Migration speed was calculated during mid-migration by measuring cluster displacement dividing by time elapsed. For delaminating clusters, delamination time was defined as the amount of time elapsed from early S9 to when the border cell cluster completely detached from the epithelium. Data was compiled, graphs generated, and statistical analysis performed using Prism (GraphPad Software).

## Acknowledgements

We thank the Westside Fly Group and Dunnwald lab for helpful discussions and the Tootle lab for helpful discussions and careful review of the manuscript. We thank Xiaobo Wang for the slbo>mCD8-GFP fly stock. Stocks obtained from the Bloomington Drosophila Stock Center (NIH P40OD018537) were used in this study. Information Technology Services – Research Services provided data storage support. This project is supported by National Institutes of Health R01GM116885. M.C.L. is partially supported by the University of Iowa Summer Graduate Fellowship and has previously been supported by the Anatomy and Cell Biology Department Graduate Fellowship.

---

**Supplemental Figure 1:**
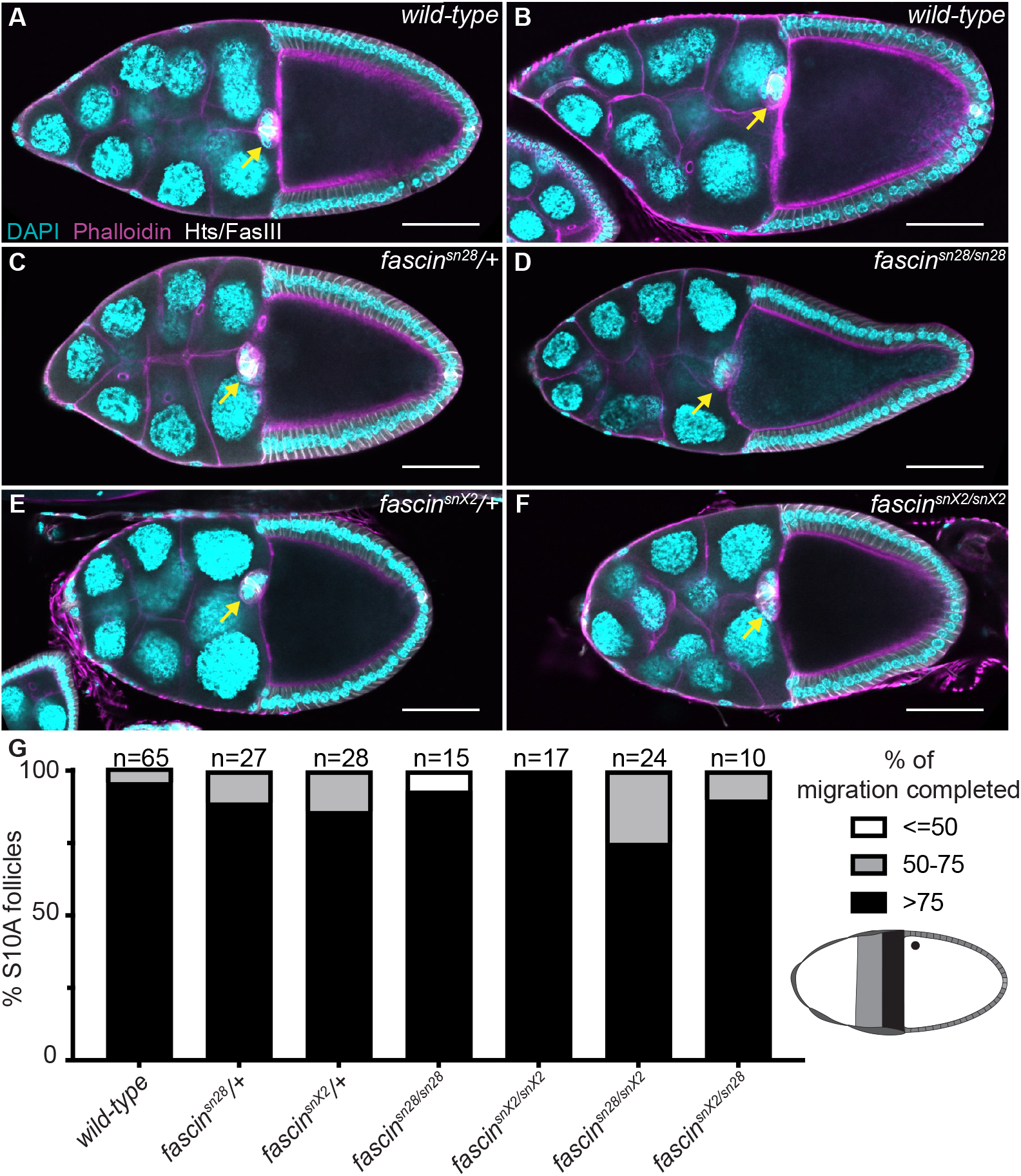
Loss of Fascin does not affect border cell migration at S10. **(A-F)** Maximum projections of 2-4 confocal slices of S10 follicles of the indicated genotypes. Merged image: Hts/FasIII (white), phalloidin (magenta), and DAPI (cyan). Scale bars = 50μm. (**A-B**) wild-type (yw). (**C**) *fascin^sn28^/+*. (**D**) *fascin^sn28/sn28^*. (**E**) *fascin^snX2^/+*. (**F**) *fascin^snX2/snX2^*. (**G**) Graph of percent of migration completed by S10A. 100% of migration completed (black), 50-75% of migration completed (grey), or less than 50% of migration completed (white). Number of follicles analyzed is indicated above the graph. Loss of Fascin by both homozygous (D, F, G) or transheterozygous *fascin* mutants (G and data not shown) do not alter the border cell cluster’s ability to reach the nurse cell-oocyte boundary by S10A compared to heterozygote *fascin* mutants (C, E, G) or wild-type controls (A-B, G).

**Supplemental Figure 2:**
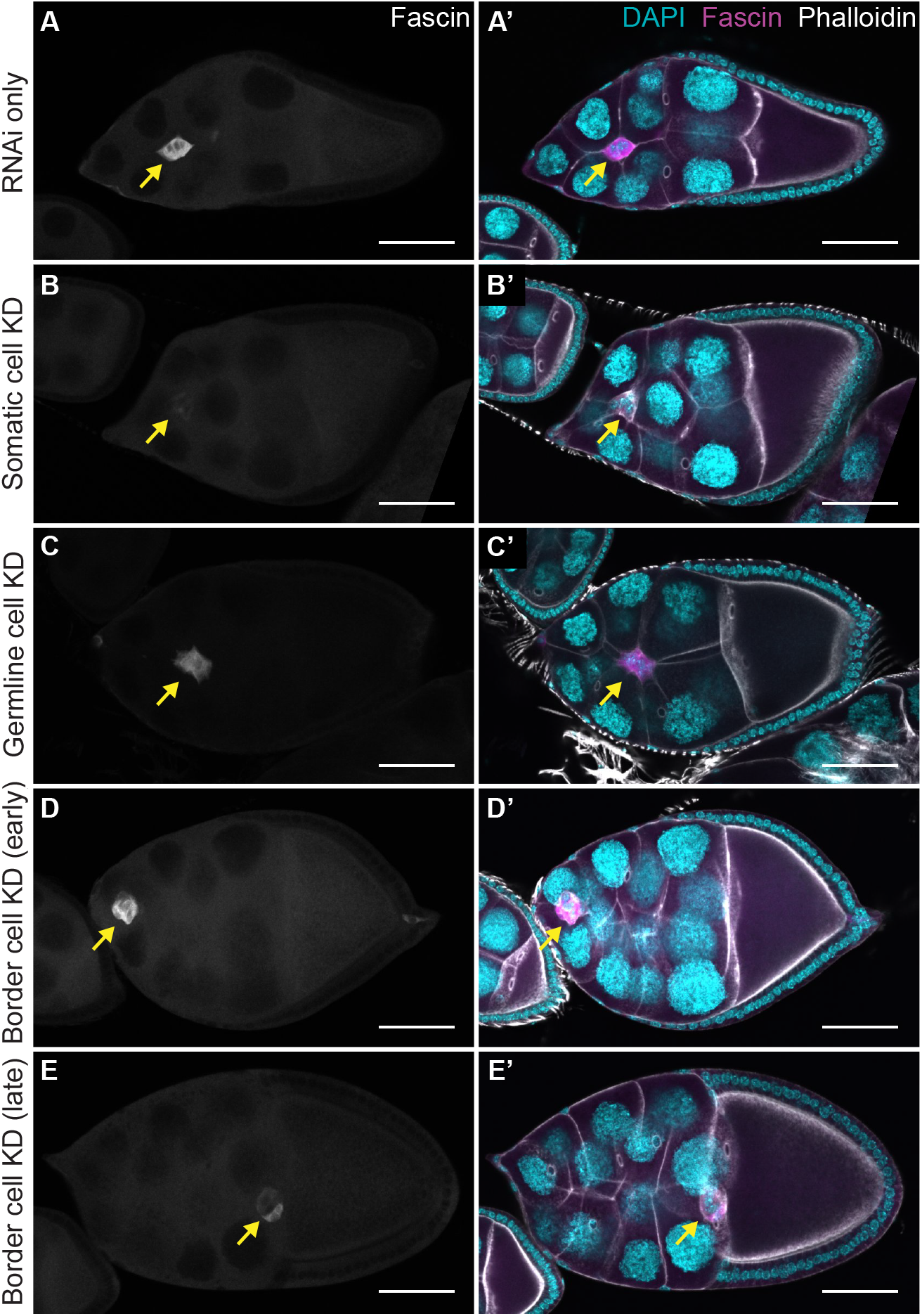
Evaluation of knockdown of Fascin with RNAi. (**A-E’**) Maximum projections of 2-4 confocal slices of S9 follicles of the indicated genotypes. (**A-E**) Fascin (white). (**A’-E’**) Merged images: Fascin (magenta), phalloidin (white), and DAPI (cyan). Yellow arrows indicate border cell cluster. Black boxes added behind text to improve text clarity. (**A-A’**) RNAi only (*+/fascin RNAi*). (**B-B’**) Somatic cell knockdown (KD) of Fascin (*c355 GAL4*/+; +/*fascin RNAi*). (**C-C’**) Germline cell KD of Fascin (*mata GAL4(3)/fascin RNAi*). (**D-D’**) Border cell KD of Fascin (*c306 GAL4*/+; +/*fascin RNAi*) at an early time point in migration. (**E-E’**) Border cell KD of Fascin (*c306 GAL4*/+; +/*fascin RNAi*) at a late time point in migration. Scale bars = 50μm. Knockdown of Fascin using the UAS/ GAL4 system is successful in the somatic cells (B-B’) and germline cells (C-C’). Knockdown of Fascin in the border cells is observed at a later point during migration (E-E’) but not at early points (D-D’).

**Supplemental Table 1.** Statistical analyses of Fascin rescue experiments for migration index quantification: (**A-C**) Table of the p-values of data represented in Figure 3G for the indicated genotypes and comparisons. (**A**) Fascin rescue experiments with the somatic GAL4 (*c355* GAL4) and all its corresponding controls. (**B**) Fascin rescue experiments with the germline GAL4 (*oskar* GAL4(2)) and all its corresponding controls. (**C**) Fascin rescue experiments with the combined somatic and germline GAL4 (*actin 5C* GAL4) and all its corresponding controls. Statistical analysis reveals that the somatic GAL4 and the combined somatic and germline GAL4 expression of GFP-Fascin in *fascin*-null mutants are significantly different compared to the *fascin*-null controls but not to the wild-type controls (A, C). The germline GAL4 expression of GFP-Fascin in *fascin*-null mutants is not significantly different from the *fascin*-null controls but is significantly different than the wild-type controls (B).

|  | *c355 GAL4/+* | *c355 GAL4/+; +/UAS-GFP-Fascin* | *fascin<sup>-/-</sup>; +/UAS-GFP-Fascin* | *c355 GAL4, fascin<sup>-/-</sup>* | *c355 GAL4, fascin<sup>-/-</sup>; +/UAS-GFP-Fascin* |
| --- | --- | --- | --- | --- | --- |
| *c355 GAL4/+* |  | >0.9999 | 0.0291 | 0.0049 | 0.9987 |
| *c355 GAL4/+; +/UAS-GFP-Fascin* |  |  | 0.0452 | 0.0096 | 0.9941 |
| *fascin<sup>-/-</sup>; +/UAS-GFP-Fascin* |  |  |  | >0.9999 | 0.0102 |
| *c355 GAL4, fascin<sup>-/-</sup>* |  |  |  |  | 0.001 |
| *c355 GAL4, fascin<sup>-/-</sup>; +/UAS-GFP-Fascin* |  |  |  |  |  |

**B**
|  | *oskar GAL4/+* | *oskar GAL4/UAS-GFP-Fascin* | *fascin<sup>-/-</sup>; +/UAS-GFP-Fascin* | *fascin<sup>-/-</sup>; oskar GAL4/+* | *fascin<sup>-/-</sup>; oskar GAL4/UAS-GFP-Fascin* |
| --- | --- | --- | --- | --- | --- |
| *oskar GAL4/+* |  | 0.9731 | 0.0092 | 0.0002 | 0.0056 |
| *oskar GAL4/UAS-GFP-Fascin* |  |  | 0.0244 | 0.0004 | 0.0146 |
| *fascin<sup>-/-</sup>; +/UAS-GFP-Fascin* |  |  |  | 0.6942 | 0.9998 |
| *fascin<sup>-/-</sup>; oskar GAL4/+* |  |  |  |  | 0.7886 |
| *fascin<sup>-/-</sup>; oskar GAL4/UAS-GFP-Fascin* |  |  |  |  |  |

**C**
|  | *actin5C GAL4/+* | *actin5C GAL4/+; +/UAS-GFP-Fascin* | *fascin<sup>-/-</sup>; +/UAS-GFP-Fascin* | *fascin<sup>-/-</sup>; actin5C GAL4* | *fascin<sup>-/-</sup>; actin5C GAL4/UAS-GFP-Fascin* |
| --- | --- | --- | --- | --- | --- |
| *actin5C GAL4/+* |  | 0.7541 | 0.0381 | <0.0001 | 0.0676 |
| *actin5C GAL4/+; +/UAS-GFP-Fascin* |  |  | 0.0012 | <0.0001 | 0.6419 |
| *fascin<sup>-/-</sup>; +/UAS-GFP-Fascin* |  |  |  | 0.175 | <0.0001 |
| *fascin<sup>-/-</sup>; actin5C GAL4* |  |  |  |  | <0.0001 |
| *fascin<sup>-/-</sup>; actin5C GAL4/UAS-GFP-Fascin* |  |  |  |  |  |

**Movie 1. Control border cell migration.** Video of S9 control follicle (*fascin^sn28^*/+*; slbo>mCD8-GFP/*+). Time listed in minutes. Images were acquired every 5.5 mins with a 20x objective. Anterior is to the right. Scale bar = 50μm. The control cluster displays single front-oriented protrusions that extend and retract throughout the migration.

**Movie 2. *fascin*-null border cell migration.** Video of S9 *fascin*-null follicle (*fascin^sn28/sn28^; slbo>mCD8-GFP/+)*. Time listed in minutes. Images were acquired every 5 mins with a 20x objective. Anterior is to the right. Scale bar = 50μm. The *fascin*-null cluster displays aberrant protrusion extensions with many protrusions extending at the same time and from the sides and back of the cluster.

**Movie 3. Control border cell delamination.** Video of early S9 control follicle (*fascin^sn28^*/+*; slbo>mCD8-GFP*/+). Time listed in minutes. Images were acquired every 5 mins with a 20x objective. Anterior is to the right. Scale bar = 50μm. The control cluster delaminates considerably faster (104min) than the *fascin*-null follicle (Movie 4).

**Movie 4. *fascin*-null border cell delamination.** Video of early S9 *fascin*-null follicle (*fascin^sn28/sn28^; slbo>mCD8-GFP/*+). Time listed in minutes. Images were acquired every 5 mins with a 20x objective. Anterior is to the right. Scale bar = 50μm. The *fascin*-null cluster delaminates significantly slower (390min) than the control follicle (Movie 3).

